# In-depth secretome analysis of *Puccinia striiformis* f. sp. *tritici* in infected wheat uncovers effector functions

**DOI:** 10.1101/2020.01.11.897884

**Authors:** Ahmet Caglar Ozketen, Ayse Andac-Ozketen, Bayantes Dagvadorj, Burak Demiralay, Mahinur S. Akkaya

**Affiliations:** Middle East Technical University, Biotechnology Program, Ankara 06800, Turkey; Division of Plant Sciences, Research School of Biology, The Australian National University, Canberra, Australian Capital Territory 2601, Australia; School of Bioengineering, Dalian University of Technology, No. 2 Linggong Road, Dalian, Liaoning 116023, China

**Keywords:** Transcriptome, secretome, effectors, yellow/stripe rust, pathogenesis, localization, function, *Puccinia striiformis* f. sp. *tritici*

## Abstract

The importance of wheat yellow rust disease, caused by *Puccinia striiformis* f. sp. *tritici* (*Pst*), has increased substantially due to the emergence of aggressive new *Pst* races in the last couple of decades. In an era of escalating human populations and climate change, it is vital to understand the infection mechanism of *Pst* in order to develop better strategies to combat wheat yellow disease. This study focuses on the identification of small secreted proteins (SSPs) and candidate-secreted effector proteins (CSEPs) that are used by the pathogen to support infection and control disease development. We generated *de novo* assembled transcriptomes of *Pst* collected from wheat fields in central Anatolia. We inoculated both susceptible and resistant seedlings with *Pst* and analyzed haustoria formation. At 10 days post-inoculation (dpi), we analyzed the transcriptomes and identified 10,550 Differentially Expressed Unigenes (DEGs), of which 6,220 were *Pst*-related. Among those *Pst*-related genes, 230 were predicted as PstSSPs. *In silico* characterization was performed using an approach combining the transcriptomic data and data mining results to provide a reliable list to narrow down the ever-expanding repertoire of predicted effectorome. The comprehensive analysis detected 14 Differentially Expressed Small-Secreted Proteins (DESSPs) that overlapped with the genes in available literature data to serve as the best CSEPs for experimental validation. One of the CSEPs was cloned and studied to test the reliability of the presented data. Biological assays show that the randomly selected CSEP, Unigene17495 (PSTG_10917), localizes in the chloroplast and is able to suppress cell death induced by INF1 in a *Nicotiana benthamiana* heterologous expression system.

## INTRODUCTION

*Puccinia striiformis* f. sp. *tritici* (*Pst*) is an obligate biotrophic fungus, which is the causative agent of wheat stripe (yellow) rust disease, a disease that can drastically reduce the production of wheat worldwide. The use of fungicides is a suitable solution for combatting *Pst*, but using resistant wheat strains to control Pst genetically would be better for the environment and more cost-effective. In order to achieve host resistance against rapid evolving virulent strains of *Pst*, understanding the molecular basis of the disease is crucial.

*Pst* infects the leaf of the wheat plant by penetrating the interior through the stomata and later differentiating into various developmental entities before finally entering into the mesophyll cells in which it can generate a feeding structure called a haustorium. The main interface for pathogen-host interactions is in the haustorium that expands between the plant cell wall and the cell membrane. This interface can activate primary host defenses when pathogen-associated molecular patterns (PAMPs) are detected by pattern recognition receptors (PRRs), resulting in PAMP-triggered immunity (PTI) [1]. Virulence is achieved by *Pst* through the deployment of secreted proteins, termed effectors, into the apoplastic fluid or inside the host [2]. The effectors are useful weapons in the fungal arsenal that allow *Pst* i) to overcome PTI, ii) to establish a suitable environment, and iii) to help proliferation [3]. Particular effectors are recognized by cytoplasmic receptor proteins in the host cell, triggering a defense response called effector-triggered immunity (ETI). Compatible and incompatible interactions depend on whether the effectors can achieve virulence by evading Resistance (R) proteins, while the R proteins evolve to detect the effectors. Therefore, evolutionary pressure imparts a driving force on the race between host resistance factors and pathogen virulence factors [4].

Investigating the secreted effectorome of the fungus guides us towards a better comprehension of the nature of pathogenic virulence and determinants that can lead to avirulence. The combination of next-generation sequencing (NGS) and data mining approaches represent a great opportunity to determine candidate effectors secreted by the fungus. Several studies attempting to construct a secretome of either *Pst* or its closest relatives (stem rust pathogen *Puccinia graminis* f. sp. *tritici* (*Pgt*) and leaf rust pathogen *Puccinia triticina* (*Ptt*) have been published. Yin and colleagues reported a set of secreted proteins generated from expressed sequence tag (EST) sequences of a cDNA library generated from *Pst* haustoria [5]. A repertoire of small-secreted proteins (SSPs) was reported using a pipeline for effector mining in the genome sequence information of *Pgt* and *Melampsora larici populina* (the pathogen that causes poplar leaf rust) [6]. The isolate PST-130 was sequenced and assembled to discover *Pst* genes and identify candidate effectors [7]. The genome sequences of *Pgt* (CRL 75-36-700-3), *Ptt* (BBDD), PST-78 (2K-041), PST-1 (3-5-79), PST-127 (08-220), PST-CYR-32 (09-001) were released by the Broad Institute (http://www.broadinstitute.org/). Cantu and colleagues used comprehensive genome analysis to identify candidate secreted effectors in two United Kingdom isolates (PST-87/7 and PST-08/21) and two United States isolates (PST-21 and PST-43) [8]. They also released data of infection-expressed genes (6 and 14 dpi) and haustorial-expressed genes (7 dpi) for PST-87/7. Another report was published using transcriptome sequencing on *Pst* isolate 104E137A to elucidate haustorial secreted proteins (HSPs) and expose the best candidate effectors [9]. A pioneer correlation analysis between genomes of seven existing races and seven new races of *Pst* was pursued to predict avirulence (Avr) determinants among SSPs [10]. In this study, 7 new races of *Pst* were sequenced with NGS and then combined with 7 older published genomes for a total of 14 races of *Pst* that were then subjected to correlation analysis to point out Avr candidates [10].

Data mining methodology is a promising strategy to identify candidate secreted effector proteins (CSEPs) in genome and transcriptome data, as the incorporation of newly generated data continually improves the prediction algorithms using deep machine learning. Likewise, our understanding of the infection and disease progression of the pathogen is steadily expanding. Therefore, new information (candidate effectors) can be generated by exhausting presently available sequence data. A combined effort of using both comparative transcriptomics and data mining to narrow down the ever-expanding sets of CSEPs will simplify and focus the work being done to discover the function of each CSEP experimentally. Only a few reports have focused on integrating both *in silico* prediction and transcriptome sequencing data on particular *Pst* races at the defined time intervals of haustoria formation and infection [8,9].

In this study, our objective is to identify differentially expressed gene sequences (DEGs) of *Pst* during the interaction with both susceptible and resistant host genotypes. We acquired *de novo* transcriptome sequencing data from both susceptible and resistant wheat cultivars infected with *Pst* for 10 days in order to discover differentially expressed secretome repertoires composed of small secreted proteins at a unique time point of the infection process after haustoria formation. Detected PstDEGs were then subjected to *in silico* prediction analysis to identify differentially expressed small secreted proteins (PstDESSPs). Then, the cataloged secretome candidates were analyzed in-depth via annotation and prediction programs to sort them into classified functions. Furthermore, a comparison of published *Pst* expression data [8,9] indicated the presence of 14 overlapping mutual CSEPs as the most promising subset for experimental validation. To test the reliability of our work, cDNA of one of the CSEP genes (Unigene17495, homologous to PSTG10917), was generated from *Pst*-infected wheat leaves (10 dpi) and cloned into *Agrobacterium tumefaciens*-compatible expression vectors. Heterologous expression of the CSEP in *Nicotiana benthamiana* revealed it was localized to the chloroplast in plant cells. Furthermore, we observed the biological function of Unigene17495 to be suppression of the INF1-mediated cell death response to *N. benthamiana*.

## MATERIALS AND METHODS

### Plant materials, growth, and pathogen inoculation

Near-isogenic wheat lines of Avocet-S and Avocet-YR10 were used to study both compatible and incompatible interactions upon *Pst* inoculation. *Pst* isolates were collected from the fields of Anatolia, Turkey and obtained from the ‘General Directorate of Agricultural Research and Policies (TAGEM) of The Ministry of Food, Agriculture, and Livestock. Avocet-S cultivars are susceptible to *Pst*TR (which has virulence towards Yr2, 6, 7, 8, 9, 25, A, EP, Vic, Mich), whereas Avocet-Yr10 is resistant. Seeds were planted in soil and grown in a growth chamber under ideal conditions (relative humidity 60%, 22°C, 16 h light, and 8 h dark). The *Pst* inoculations were performed when the seedlings reached the two-leaf stage by spraying freshly collected urediniospores directly onto the leaves. Before the inoculation, urediniospores (100 mg) were germinated for 5 min at 42°C and treated with mineral oil (1 mL) to enhance their attachment to the leaf blade. For the control samples (mock inoculations), only mineral oil was used. After air-drying for 20 min, the plants were kept in the dark at 10°C overnight in the presence of continuous mist in order for infection to take place. After overnight infection, the standard growth conditions were resumed. The wheat leaves were collected at 10 days post-inoculation (10 dpi) when the disease symptoms were visible on the susceptible cultivar.

### RNA extraction, *de novo* transcriptome sequencing, assembly, and annotation

The flow chart in Fig 1 describes the overall protocol followed for collecting infected leaf samples using homogenization with liquid nitrogen and a sterile mortar and pestle. For 0.1 g leaf sample, 0.02 g Poly(vinylpolypyrrolidone) (PVPP) was added during mortar and pestle grinding. For each condition, three biological replicates were homogenized together. Total RNA isolation was carried out using QIAzol Lysis Reagent (Qiagen) by following the manufacturer’s protocol. A NanoDrop was used for the spectrophotometric quantification of the total RNA samples. The samples were shipped to Beijing Genomics Institute (BGI) in absolute ethanol for sequencing. BGI measured the RNA quality of the samples on an Agilent 2100 Bioanalyzer, and the samples with a RIN value above 8 were allowed to proceed to mRNA enrichment and cDNA synthesis using random hexamers. Single nucleotide adenine was added to the purified and end-repaired short cDNA fragments. After adapter ligation, size selection was used to obtain templates for PCR amplification. The Illumina HiSeq™ 2000 platform was used to perform *de novo* transcriptome sequencing. BGI-Shenzhen (Shenzhen, China) conducted all of the protocol steps from mRNA enrichment to sequencing. All quality controls were evaluated during sequencing using the Agilent 2100 Bioanalyzer and ABI StepOnePlus Real-Time PCR System for quantification and qualification of the sample libraries.

**Fig 1.**
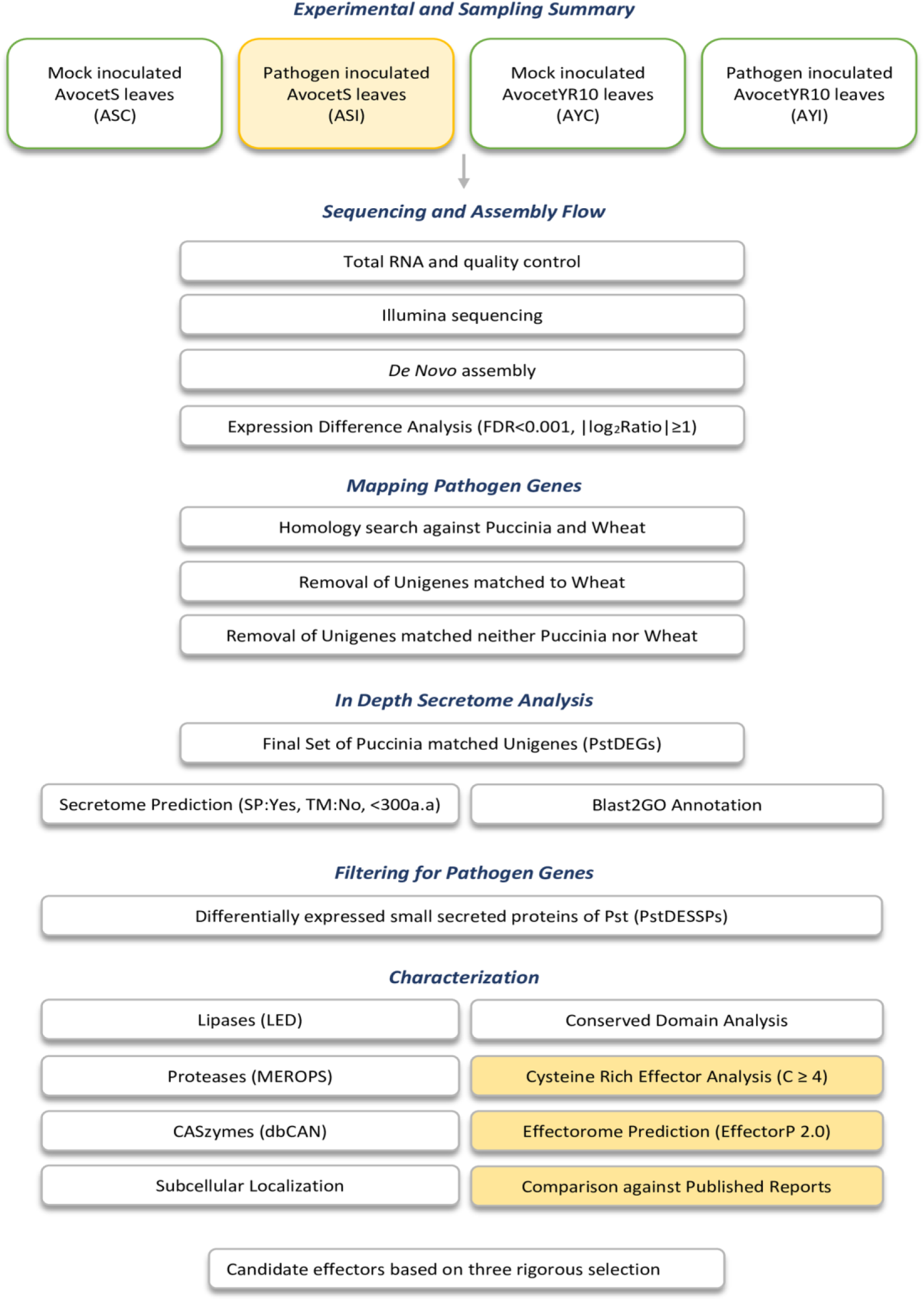
Flowchart to summarize sampling, sequencing, and *in silico* analysis. Pipeline for the steps performed to achieve *de novo* assembly, expression analysis, and mapping of *Pst* genes. * = FDR (False Discovery Rate).

Before assembly, clean reads were obtained from raw reads using internal BGI software (filter_fq) that follows three main parameters: (1) elimination of reads with adaptors, (2) removal of reads with unidentified nucleotides larger than 5%, (3) elimination of low-quality reads (if the percentage of the low-quality value (Q ≤ 10) reads are higher than 20%, they are removed). For *de novo* transcriptome assembly, clean reads of each library were assembled into contigs and unigenes by the Trinity program (release-20130225, http://trinityrnaseq.sourceforge.net/) [11].

Unigenes were aligned using the Blastx program (e-value < 0.00001) and the following protein databases: NR (NCBI, non-redundant database) [12], Swiss-Prot (EMBL protein database) [13], KEGG (Kyoto Encyclopedia of Genes and Genomes) [14], and COG (Clusters of Orthologous Groups) [15]. The direction of the unigene sequences was determined using the best alignment scores. NR, Swiss-Prot, KEGG, and COG were sourced to predict the coding region (CDS) of unigenes in the subsequent order of priority if a conflict in sequence orientation arose. ESTScan software was performed on unigenes with no successive alignments [16]. The results for NR, NT, Swiss-Prot, KEGG, and COG database alignments were also used for unigene function annotation. To obtain gene ontology (GO) annotation from NR annotations, the Blast2GO program (v2.5.0) was performed (e-value < 0.00001) on fungi databases (taxa, 4751) [17]. Web Gene Ontology Annotation Plot (WEGO) software was used for the classification of GO annotations [18]. The KEGG database was employed to investigate the metabolic pathway analysis of unigenes [14].

### Expression difference analysis

Differentially expressed genes (DEGs) were identified using the FPKM method also known as RPKM method [19]. In this method, FPKM stands for fragments per kb per million reads, and RPKM stands for reads per kb per million reads. We used the following calculation formula:

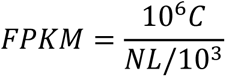

FPKM signifies the expression of Unigene A, C is the number of fragments that uniquely aligned to Unigene A, N is the total number of fragments that uniquely aligned to all unigenes, and L is the base number in the CDS of Unigene A. As we obtained FPKM values for each unigene, we calculated FPKM ratios between two samples at a time. We defined differentially expressed genes (DEGs) to have a FPKM ratio ≥ 2 and a false discovery rate (FDR) ≤ 0.001. We executed GO functional analysis (*P*-value ≤ 0.05), including GO functional enrichment and functional classification on DEGs. Furthermore, we carried out the KEGG pathway analysis (Q-value ≤ 0.05) for pathway enrichment to further understand biological functions [20].

### Identification of pathogen-associated DEGs

We identified our DEGs through transcriptome data obtained from pathogen-inoculated wheat leaves. However, we needed to eliminate any possible contaminating sequences from the host. To accomplish this, we constructed a local database (named ‘The Pucciniale database’) by merging protein data of *Puccinia striiformis* f. sp. *tritici* (isolate 2K41-Yr9, race PST-78), *Puccinia graminis* f. sp. *tritici* (strain CRL 75-36-700-3, race SCCL), and *Puccinia triticina* (isolate 1-1, race 1-bbbd). All protein data was obtained from the Broad Institute Puccinia website (http://www.broadinstitute.org/). We also merged protein data of the WheatD genome of *Aegilops tauschii* (wheatD_final_43150.gff.cds, downloaded from GIGA_DB ‘http://gigadb.org/’) [21] to ‘The Pucciniale database’. We performed a BlastX analysis (e-value cut-off ≤ e^−10^) on DEGs using our local database of Pucciniale and wheat [22]. If the hit matched with the wheat proteome or even if it produced a significant hit with the Pucciniale proteome, we discarded the sequences following the decision rule. The best hits within the Pucciniale proteome were evaluated as PstDEGs. The sequences with no significant hits against neither the pathogen nor host were assessed as unknown novel DEGs.

### Construction and characterization of secretome data

We predicted the ORFs and protein sequences from the PstDEG data using the ORFPredictor web tool (http://bioinformatics.ysu.edu/tools/OrfPredictor.html). We followed a widely accepted secretome prediction pipeline to construct our own differentially expressed secretome database [6,23,24]. The predictions were made following three criteria: 1) presence of N-terminus signal peptide region, 2) absence of transmembrane helices, and 3) mature protein length should be smaller than 300 amino acids. SignalP (version 4.1) was used to predict the presence of a secretion signal [25]. We estimated transmembrane helices using the TMHMM webtool (version 2.0, http://www.cbs.dtu.dk/services/TMHMM/) [26]. Filtered sequences were termed as differentially expressed small-secreted proteins of *Puccinia striiformis* f. sp. *tritici* (PstDESSPs). The same filtering pipeline was applied to total proteins obtained from the ‘Broad Institute database’ for *Pst* and *Pst* relatives (*Pgt* and *Ptt*) (http://www.broadinstitute.org/). Four secretome data sets (PstDESSPs, PstSSPs, PgtSSPs, PttSSPs) were subjected to effector prediction via the EffectorP 1.0 and 2.0 programs (http://effectorp.csiro.au/) [27,28]. The Blastp program was used (E-value < e^−5^) for characterization of PstDESSPs by scanning against the following databases: the lipase engineering database (http://www.led.uni-stuttgart.de/), the protease and protease inhibitors (MEROPs) database (http://merops.sanger.ac.uk/) [29], the carbohydrate-active enzymes (CAZymes) database (http://csbl.bmb.uga.edu/dbCAN/) [30], the fungal peroxidase database (fPoxDB, http://peroxidase.riceblast.snu.ac.kr/) [31], and the pathogen-host interaction database (http://phi-blast.phi-base.org/) [32]. The ClustalW program was executed for multiple alignment analysis of PstDESSPs (https://www.ebi.ac.uk/Tools/msa/clustalo/) [33]. The iTOL web-tool was used to construct a phylogenetic tree using multiple alignment results (https://itol.embl.de/) [34]. Small secreted cysteine-rich effector proteins were assessed as cysteine-rich if the protein possessed four cysteine residues or more. A comparison between infection- and haustoria-expressed genes was done using a similarity search using Blastp (e-value cut-off ≤ e^−10^) on publicly available data [8,9]. We scanned for PstDESSPs overlapping with *Avr* determinant candidates originally interpreted using correlation analysis by Xia and colleagues [10]. Subcellular localization predictions were done on mature proteins (without N-terminus signal peptides) to forecast translocation sites in the host plant using four different tools: TargetP 1.1 (http://www.cbs.dtu.dk/services/TargetP/) [35], Localizer (http://localizer.csiro.au/) [36], WolfPSORT (https://wolfpsort.hgc.jp/) [37], and ApoplastP (http://apoplastp.csiro.au/) [38]. All of the analyses, annotations, and predictions regarding PstDESSPs are displayed in Table S3.

### Cloning of Unigene17495 (Pstg10917)

The cloning primers were designed using the Pstg10917 sequence with an N-terminus CACC site in forward primers (with and without signal peptide (SP)) and no stop codon in reverse primers (Table S4). Total RNA samples of infected Avocet-S leaves (10 dpi) used in transcriptome sequencing were sourced to synthesize cDNA after DNase treatment. The Transcriptor First Strand cDNA Synthesis Kit (Roche) was used following the manufacturer’s protocol, except that we preferred using a mixture of both random hexamer primers and oligo(dT) primers at equal volumes to the reaction primer. The effector of interest, with and without SP, was amplified from cDNA, cloned into the pENTR/D-TOPO vector (Invitrogen), and then subsequently cloned into the pK7FWG2 expression vector [39] utilizing Gateway Cloning. Both colony PCR and sequencing analysis validated the cloning of constructs at the proper positions. Then, the plasmid constructs were electroporated into *Agrobacterium tumefaciens* GV3101.

#### Agrobacterium tumefaciens-mediated gene transfer

*A. tumefaciens* infiltration experiments were performed using the previously described protocol [40] with slight alterations. *A. tumefaciens* GV3101 strains carrying expression vectors were plated on selective agar media. After two days, bacteria were scratched and suspended in sterile water. Bacteria were collected by centrifugation at 4,000 rpm for 4 min at room temperature (24-25°C). Bacteria were washed two times with sterile water and two times with *A. tumefaciens*-mediated gene transfer-induction medium (10 mM MES, pH 5.6, 10 mM MgCl_2_). The final cell concentration was adjusted to 0.4 at A_600nm_ for infiltration unless otherwise stated. The inoculums were injected into *Nicotiana benthamiana* leaves using a needleless syringe. The expressions of the gene of interests were observed in the next 2-4 d.

### Confocal microscopy

*A. tumefaciens* GV3101 cells carrying pK7FWG2/Pstg_10917 and pK7FWG2/ΔSP-Pstg_10917 were injected into *N. benthamiana* leaves (4-5 weeks old) for transient expression. After 2-3 d, leaves were cut into small pieces of 3-5 mm length and soaked in tap water. A Leica 385 TCS SP5 confocal microscope (Leica Microsystems, Germany) was used for subcellular localization imaging. The wavelengths used for imaging GFP fluorescence were excitation at 488 nm and emission at 495-550 nm. Alternatively, the excitation and emission wavelengths for chloroplast autofluorescence were in the far-infrared (>800 nm).

### Cell death suppression assay

The pTRBO/GFP plasmid for GFP expression (courtesy of the Kamoun Lab, Sainsbury Laboratory, Norwich, UK) and the pK7FWG2/SP-GFP plasmid [41] for SP-GFP were used as negative controls in the cell death suppression assays. The *A. tumefaciens* GV3101 cells carrying the negative control plasmids (GFP and SP-GFP) or the effector of interest (with and without SP) were injected into a *N. benthamiana* leaf. After 24 h, *A. tumefaciens* GV3101 carrying cell death elicitor pGR106/Inf1 (a gift from the Bozkurt lab, Imperial College, London, UK) was injected in the same spot without damaging leaf tissue. Cell death was measured at 4 dpi.

## RESULTS AND DISCUSSION

### Sequencing of *Pst*-infected wheat leaves and *de novo* transcriptome assembly

We have collected total RNA from *Pst*-infected wheat leaves of susceptible and resistant seedlings to analyze gene expression differences between the compatible and incompatible pathogen-host interactions. The sequence reads have been deposited in the NCBI BioProject database under the accession number PRJNA545454. A total of 9,204,786 clean reads for *Pst*-infected susceptible wheat (Avocet-S_PST), and 6,899,596 clean reads for *Pst*-infected resistant wheat cultivar (Avocet-YR10_PST) were obtained after quality filtering. Their Q20 values were 98.16% and 97.90%, respectively (Table 1).

**Table 1.**
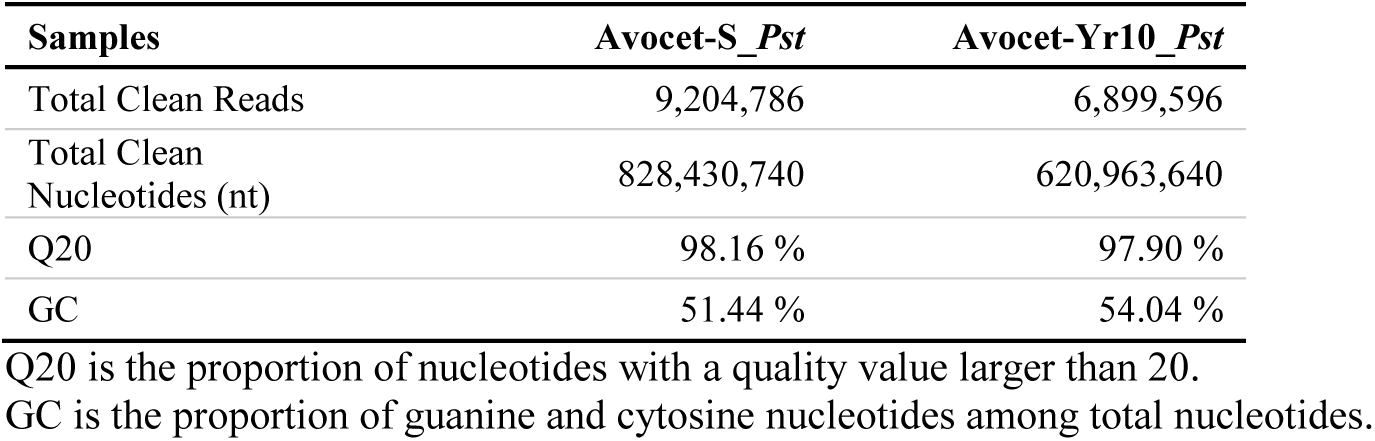
Sequencing statistics for *Pst* inoculated/infected wheat transcriptome.

The clean reads were assembled into more than 100,000 contigs with an average length of greater than 200 nucleotides using Trinity software (Table 2). Unigenes were clustered based on their level of similarity with each other. A cluster was defined as containing unigenes with more than 70% similarity. We generated 12,956 distinct unigene clusters for Avocet-YR10_PST and 13,564 for Avocet-S_PST (Table 2). A total of 42,275 and 55,004 unigenes came from single genes within the two sets, respectively. This data is represented in Table 2 as the distinct singleton. The length distribution of assembled contigs and unigenes is provided in Fig S1.

**Table 2.** *De novo* assembly statistics for *Pst* inoculated/infected wheat transcriptome.

|  | Samples | Number | Length (nt) | Mean Length (nt) | N50 | Distinct Clusters | Distinct Singleton |
| --- | --- | --- | --- | --- | --- | --- | --- |
| Contig | Avocet-Yr10_*Pst* | 105,866 | 23,189,064 | 219 | 249 | - | - |
|  | Avocet-S_*Pst* | 128,680 | 29,848,685 | 232 | 272 | - | - |
| Unigene | Avocet-Yr10_*Pst* | 55,231 | 18,753,197 | 340 | 371 | 12,956 | 42,275 |
|  | Avocet-S_*Pst* | 68,568 | 24,329,132 | 355 | 394 | 13,564 | 55,004 |
|  | All | 61,105 | 26,978,874 | 442 | 490 | 15,016 | 46,089 |

Distinct clusters are similar (more than 70%) unigenes, and these unigenes may originate from the same gene or homologous gene; whereas distinct singletons represent the unigene come from a single gene.

### Functional annotation and classification of *de novo* transcriptome

The unigenes obtained from both Avocet-S_PST and Avocet-YR10_PST were merged into a single integrated data set to conduct an overall interpretation of transcriptome sequencing. The unigenes were aligned against the following databases: NR, NT, Swiss-Prot, KEGG, COG, and GO for annotation via the Blastx program (e-value < 0.00001). Overall, 26,843 of 61,105 unigenes, corresponding to 43.9%, were annotated in one or more databases (Table 3). The detailed annotation results are listed in Table S1. The NR database provided the highest number of annotations. Approximately half of the annotated unigenes shared high similarity (>60%) and aligned to either *Pst* genes (41.4%) or genes of other fungal species (Fig 2). The results of COG classification and GO functional annotation of the unigenes are presented in Fig S2.

**Fig 2.**
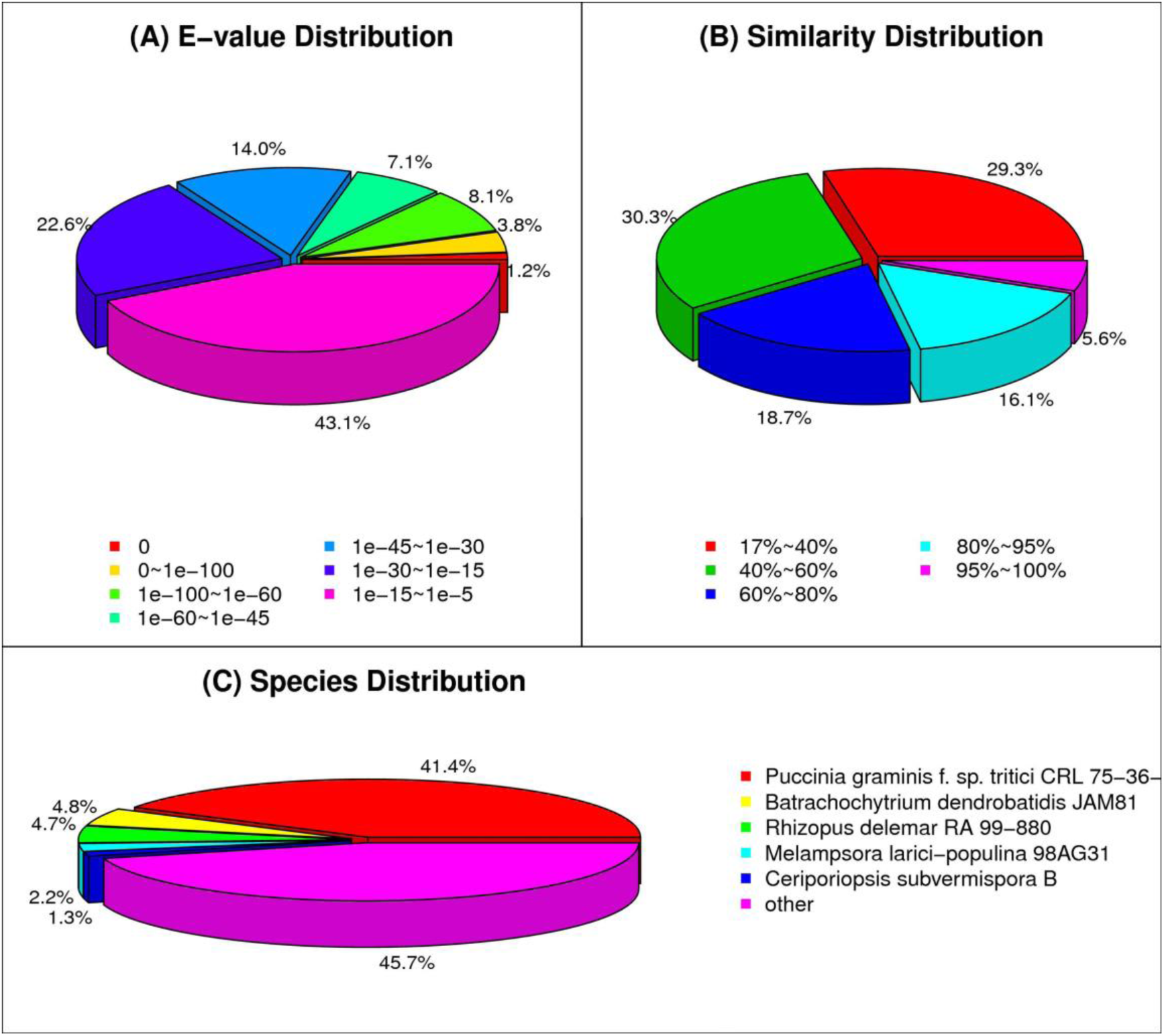
Statistics of the NR classification of unigenes. (A) The e-value distribution of the alignment results of NR annotation. (B) The similarity distribution and (C) the species distribution of the results of NR annotation.

**Table 3.** Annotation statistics of the assembled unigenes of the transcriptome sequencing project.

| Sequence file | NR | NT | Swiss-Prot | KEGG | COG | GO | All |
| --- | --- | --- | --- | --- | --- | --- | --- |
| All-Unigene.fa | 24,933 | 9,945 | 16,318 | 16,275 | 13,278 | 10,015 | 26,843 |

### Discovery of differentially expressed unigenes (DEGs)

We used the FRKM (RPKM) method [19] to determine DEGs between compatible and incompatible interactions. We identified 10,550 DEGs with a false discovery rate (FDR) of less than 0.001, and an expression difference of at least two-fold (Fig 3).

**Fig 3.**
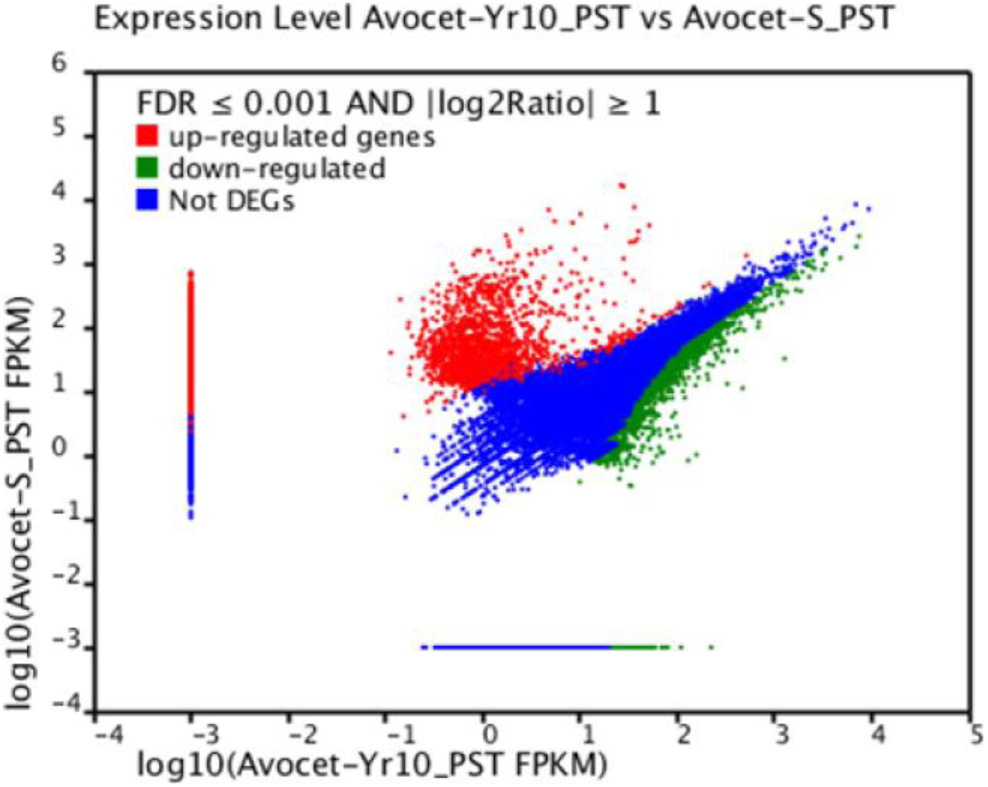
Distribution of differentially expressed unigenes (DEGs). Red: Up-regulated genes (6851), green: downregulated genes (3699), and blue: no differential expression observed with the parameters FDR ≤ 0.001 and | log2ratio | ≥ 1.

We mapped the DEGs to our custom database constructed by merging the pathogen and host genomes. Local Blastx analysis (e-value < 0.00001) revealed that 6,220 of the DEGs aligned best to the *Pucciniales* genome and 2,880 of the DEGs aligned best to the wheat genome. The remaining DEGs yielded no significant homology to either the pathogen or the host; hence, they are considered to be unknown novel DEGs. The 6,220 *Pst*-mapped DEGs (PstDEGs) were selected as the differentially expressed, pathogen-associated candidate genes to proceed with for secretome analysis.

The identified PstDEGs were subjected to Blast2GO analysis to elucidate their functional classification (Table S2). A graphical view of the molecular functions and biological processes of the PstDEGs is displayed in Fig 4. We were able to annotate the majority (75.8%) of PstDEGs with their corresponding functional classification. The distribution of molecular function between proteins with catalytic activity (61.73%) and binding activity (61.07%) attributes were practically equal in the Blast2GO level 2 analysis (Gene Ontology (GO) specificity increases as the “level” of the analysis gets higher). During level 3 analysis of the PstDEGs, we observed that the proteins with catalytic activity belonged to the hydrolase, transferase, and oxidoreductase categories (Fig 4A). The Blast2GO analysis illustrates that the pathogen undergoes radical changes during disease progression. A large number of PstDEGs (6,220 genes compared to the total predicted protein number of *Pst* genes which is over 20,000) were sorted into various categories of biological processes (Fig 4B). Therefore, we wanted to determine the number of PstDEGs that sorted as enzymes. In total, we cataloged 1,824 PstDEGs as enzymes. Among them, hydrolases were the most common, followed by transferases and oxidoreductases (Fig 4C).

**Fig 4.**
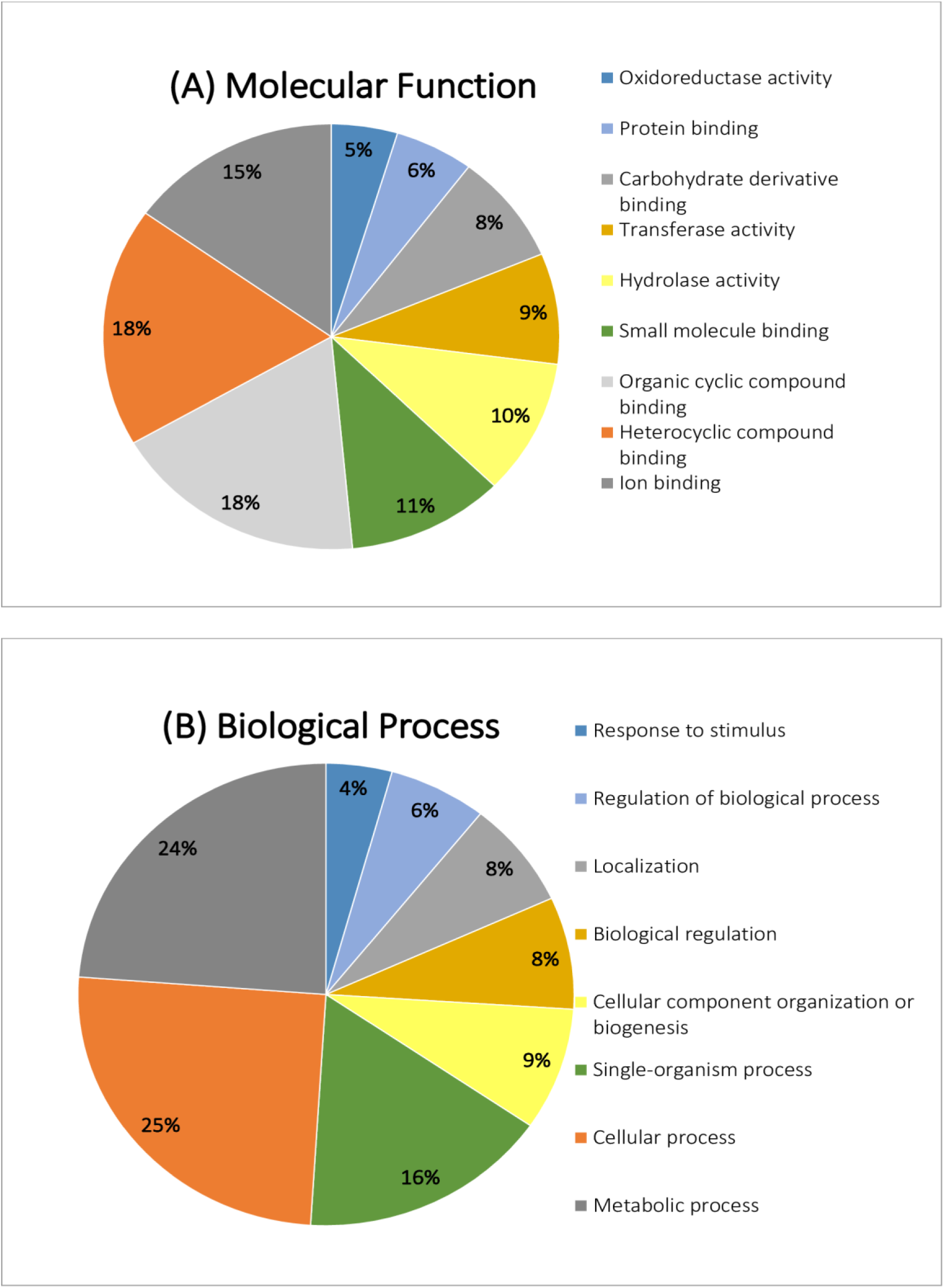
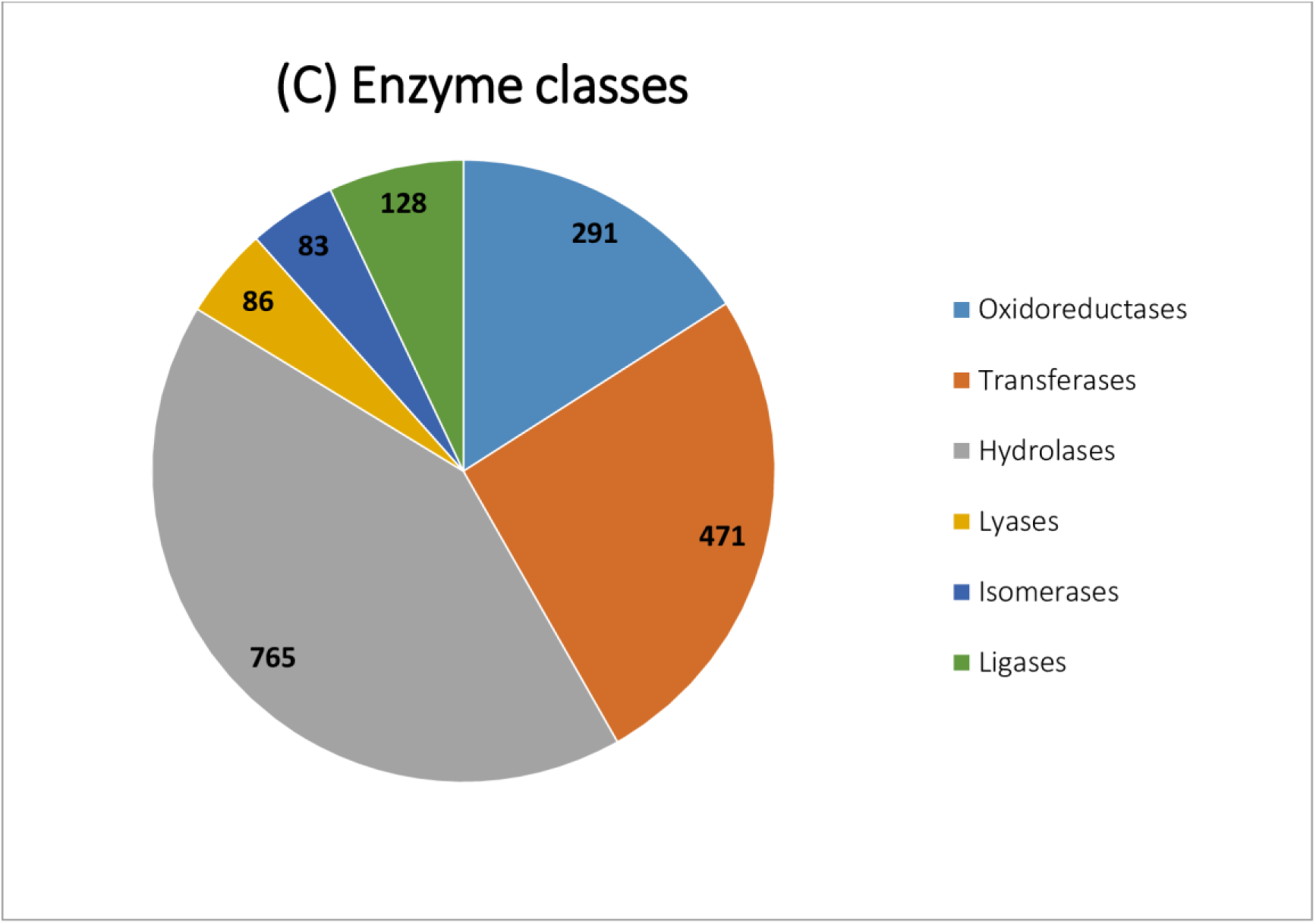
Functional annotations of PstDEGs via the Blast2Go program. **(A)** Molecular function analysis (Level 3) of PstDEGs. **(B)** Biological process classification (Level 2) of PstDEGs. **(C)** Enzyme class distribution of PstDEGs.

### Identification and characterization of the differentially expressed secretome

We made use of a previously constructed pipeline defined and validated by several research groups (6, 23, 24). We extracted the secretome repertoire of 230 small-secreted proteins of *Puccinia striiformis*, which are referred to as PstDESSPs, by selecting all of the differentially expressed genes identified in our transcriptome data (6,220 differentially expressed genes (DEGs)). We identified other small-secreted proteins by applying the same pipeline on all of the available genome sequences of *Puccinia* species: *Pst*, *Pgt*, and *Ptt* (http://www.broadinstitute.org/). We then compared all of the identified secretomes in a method similar to that used in the study published by Xia *et al*., 2017 [10]. Among 230 small secreted proteins, 94 were predicted as effectors using ‘Effector 2.0’ software, and were then classified as Pst-Small Secreted Candidate Effectors (PstSSCEs) (Fig 5A). The percentage of PstDESSPs within the data set of all differentially expressed genes was 3.7% (230 out of 6,220). This ratio appeared significantly smaller when compared to the hypothetical prediction of small secreted proteins within PST-78 (6.5%, 1,332 out of 20,482) and the other *Pucciniale* genomes. Therefore, we concluded that the number of actively involved small secreted proteins at 10 dpi is indeed much lower than what was hypothetically expected. This indicates that the hypothetical number of PstDESSPs can be exaggerated when obtained using only transcriptome predictions from genome sequences.

**Fig 5.**
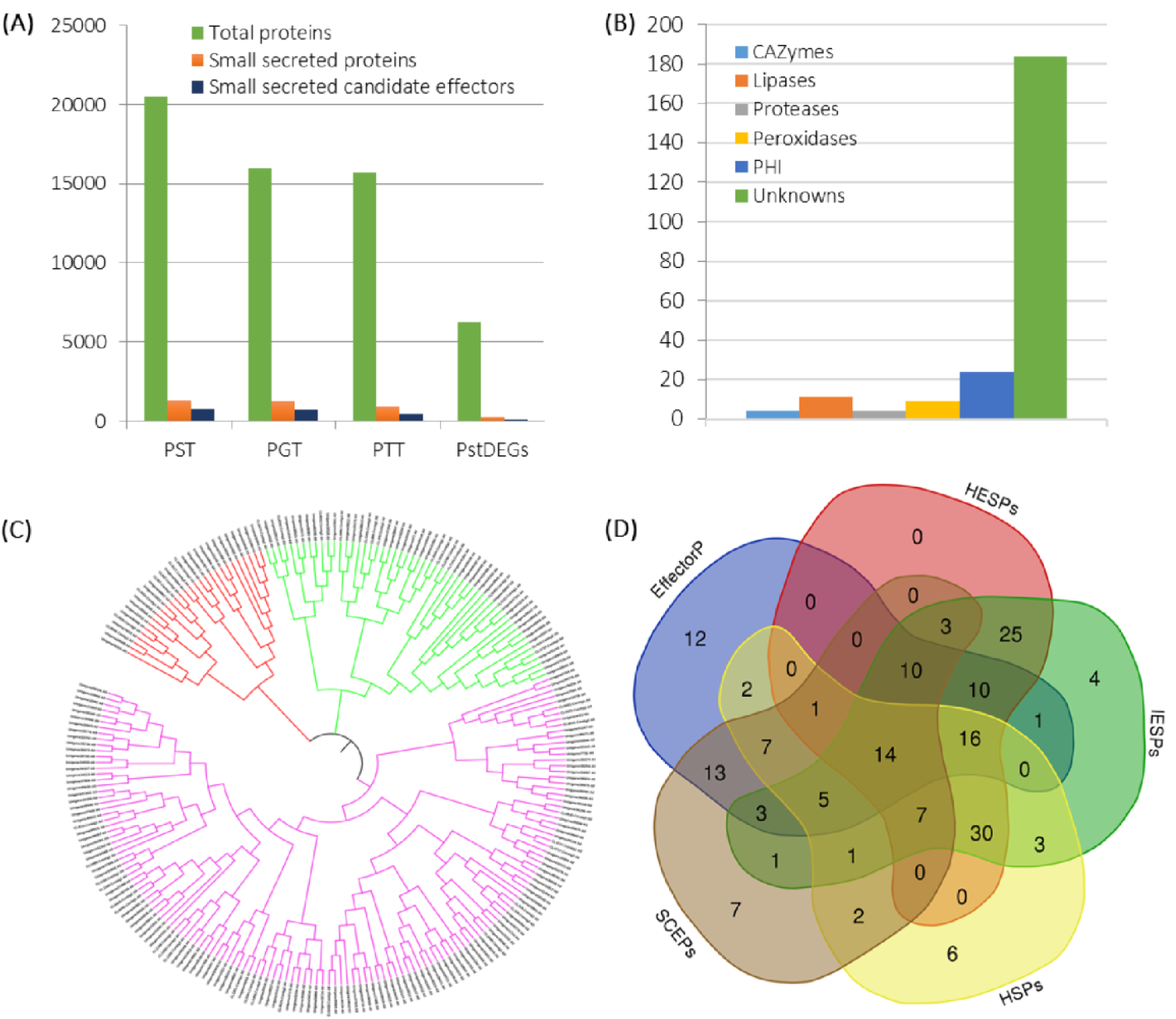
Characterization of the PstDESSPs. (A) The number of total proteins, small secreted proteins, and small secreted candidate effectors among our data (PstDEGs) and three wheat rust fungi genomes: *Pst, Pgt* (*P. graminis* f. sp. *tritici*), and *Ptt* (*P. triticina*). (B) Distribution of PstDESSPs for functional annotations. (C) Phylogenetic analysis based on the protein sequence similarity of PstDESSPs. (D) Characterization and comparison of PstDESSPs in five categories: small cysteine-rich effector proteins (SCEPs), predicted to be an effector candidate by a machine learning algorithm (EffectorP 2.0-positive), and consistent with previous sequencing reports in the literature of candidate rust effectors in haustorial (HESPs) or infection (IESPs) expressed secreted proteins [8], and haustorial secreted proteins (HSPs) [9].

Certain types of gene functions are useful for the fungus in pathogen-host interactions, including Carbohydrate-Active Enzymes (CAZymes), proteases, peroxidases, and lipases. The PstDESSPs were compared against several distinguished public databases (MEROPs, dbCAN, LED, fPoxDB) in order to predict their functions. We identified 4 proteases and protease inhibitors, 4 CAZymes, 11 lipases, and 9 peroxidases among the PstDESSPs (Fig 5B). We searched the Pathogen-Host Interaction (PHI) database to predict the virulence attribute of each PstDESSP based on their sequence similarity. A total of 24 of the PstDESSPs exhibited significant similarity to at least one entry in the PHI database. For example, Unigene31938 (Pstg04010) had a predicted ‘copper-zinc superoxide dismutase’ domain match with PHI383 (*C. albicans*) and PHI6412 (*P. striiformis*) (Table S3). Although the level of similarity barely exceeded the cutoff value, both of these entries have a superoxide dismutase domain. PHI6412 (PsSOD1) has been reported to increase the resistance of the pathogen against host-derived oxidative stress [42]. The results suggest that all of the PstDESSPs that matched with entries in the PHI database are worthy enough of further exploration employing *in vivo* systems. Hence, a comparison against multiple databases provides a better way to discover CSEPs and understand the importance of the PstDESSPs in Table S3. Furthermore, a considerable percentage (80%) of the PstDESSPs remain uncharacterized (Fig 5B), due to the tendency of the candidate effectors to only show homology to the pathogenicity related domains and not to other typical functional domains.

To understand the relationship between PstDESSPs, we generated a phylogenetic tree using an analysis of multiple sequence alignments. The outcome suggests that there are no significant conserved sequences among the 230 PstDESSPs. In general, we identified three main branches (or classes) of PstDESSPs based on sequence similarities. The purple branch in Fig 5C possesses a substantial amount of the PstDESSPs and is also subdivided into two smaller arms. No significant conserved motifs were detected among the PstDESSPs, except for the Y/F/WxC motif (Table S3), which is common to the powdery mildew [43].

The PstDESSPs determined in this study were compared with other published expression data sets of *Pst* to investigate whether the SSPs in this study were unique or whether they were similar to haustorial differentially expressed effector candidates. Cantu and colleagues studied the expression data for effectors expressed in infected leaf and haustoria samples and provided a pioneer list of SSP tribes in their analysis (7, 8). We analyzed the similarity between tribes of SSPs via the Blastp program (e-value < 0.00001). A total of 116 PstDESSPs were found to share significant homology to haustoria-expressed secreted effector proteins (HESPs), and 133 of PstDESSPs were found to be homologous to infected leaves-expressed secreted effector proteins (IESPs) (Table S3). The remaining 96 PstDESSPs that did not match with HESPs or IESPs were found to be differentially expressed at 10 dpi in this study. Comparison with the list of haustorial secreted proteins (HSPs) published by Garnica *et al.* indicated that 94 of the PstDESSPs are drastically similar to HSPs [9]. Overall, the number of PstDESSPs that we identified that are not similar to any of those in previously reported data sets is 32, signifying their exclusivity to the present study. Xia and colleagues scrutinized the effector candidates for avirulence (Avr) by comparing the expression data sets of Cantu *et al*., 2013 [8], [10]. We compared our list of PstDESSPs to the list of Avr candidates and the results suggest that three previously overlooked Avr candidates and six known Avr candidates are present among our PstDESSP data set (Table S5).

The cysteine content of a candidate effector protein has long been a determinant for making effector predictions [8–10,23,24]. Although several effectors have been discovered to be not rich in their cysteine composition, such as AvrM [44] and AvrL567 [45], we examined the PstDESSPs in terms of cysteine possession. We identified 64 cysteine-rich small-secreted effector proteins (SCEPs) that have four or more cysteine amino acid residues in their mature protein length (not including the N-terminal signal peptide) (Table S5).

To find the most promising subset of candidate effectors, we focused on the PstDESSPs that produced positive results in the EffectorP analysis, had cysteine residue counts of more than three, and had homologous matches within the previously mentioned reports (HESPs, IESPs, and HSPs). A total of 14 PstDESSPs met all five criteria and were defined as reliable CSEPs for further investigations (Fig 5D). A more detailed description of the features of the 14 CSEPs is provided in Table 4.

**Table 4:** List of the most promising CSEPs overlapped in all five classes in Figure 5D.

| ID | CSEP Properties |  |  | Subcellular Localization Prediction |  |  |  |  |  |
| --- | --- | --- | --- | --- | --- | --- | --- | --- | --- |
|  | Corresponding Pst78 Homologs | Conserved Domains | Length (a.a.)<sup>a</sup> | Cys<sup>b</sup> | DE<sup>c</sup> | TP<sup>d</sup> | Loc.<sup>e</sup> | WolfPSORT | ApoplastP |
| **CL887.Contig1** | PSTG_16009 |  | 138 | 10 | Up | - | - | C | AP |
| **CL887.Contig2** | PSTG_16009 |  | 138 | 10 | Up | - | - | C | AP |
| **Unigene13023** | PSTG_03450 |  | 100 | 10 | Up | - | - | C | AP |
| **Unigene17495** | PSTG_10917 |  | 111 | 6 | Up | C | C | M | AP |
| **Unigene28760** | PSTG_14090 | DPBB_1 superfamily | 112 | 6 | Up | - | - | C | AP |
| **Unigene29179** | PSTG_04274 |  | 82 | 6 | Up | - | - | C | AP |
| **Unigene31294** | PSTG_00994 |  | 254 | 10 | Up | - | N | N | NonAP |
| **CL5364.Contig2** | PSTG_01880 |  | 205 | 6 | Up | - | - | ER | NonAP |
| **CL5364.Contig3** | PSTG_01881 |  | 205 | 6 | Up | - | - | N | AP |
| **Unigene33801** | PSTG_05079 |  | 92 | 6 | Up | - | - | C | AP |
| **Unigene36354** | PSTG_08969 |  | 196 | 8 | Up | C | - | C | AP |
| **Unigene36901** | PSTG_03879, partial |  | 49 | 6 | Up | - | - | N | AP |
| **Unigene38807** | PSTG_14378 |  | 191 | 10 | Up | - | - | G | AP |
| **Unigene7930** | PSTG_02173 |  | 109 | 4 | Up | - | - | C | AP |

<sup>a</sup>Length represents the length of mature proteins without N-terminus signal peptide part.
<sup>b</sup>Cysteine residue number in the mature protein.
<sup>c</sup>Differential expression compatible interaction over incompatible interaction.
<sup>d</sup> The results of TargetP (TP) program,<sup>e</sup> Localizer (Loc.) Program. Abbreviations are C: Chloroplast, M: mitochondria, N: nucleus, G: Golgi, E.R.: endoplasmic reticulum, Cy: cytosolic, AP: apoplastic, NonAP: non-apoplastic. All predictions were performed using mature protein sequences under the assumption that mature proteins translocate into the host cell.

### Subcellular localization of Unigene17495 (Pstg_10917)

One of the CSEPs from Table 4 was chosen at random to test the reliability of the presented PstDESSPs data. Unigene17495 (Pstg_10917) was predicted to have a transit peptide sequence following the N-terminus secretion signal. Thus, if the effector candidate succeeds in translocation within the host cell, it may be able to partially target the chloroplast, the site of control for programmed cell death (PCD) against both biotic and abiotic stressors. The heterologous expression of Unigene17495 with a GFP tag on *N. benthamiana* leaves showed chloroplast localization (Fig 6). Its localization to the chloroplast despite being predicted for secretion outside the cell was unexpected. However, re-checking the predictions by other algorithms; ‘TargetP’ in addition to predicting a secretion signal, only after its removal, it also detected a transit peptide within the remainder sequence (111 a.a.). On the other hand, ‘Localizer’ analysis predicted a transit peptide overlapping the secretion signal region. Therefore, the experimentally shown chloroplast localization, instead of originally predicted secretion outside of the cell, can be explained by the presence of a transit peptide and a possible disruption of the secretion signal due to a particular condition in the vicinity of the molecule. Alternatively, in the natural host, trafficking in both locations may be possible or there is a slight change in the physiological condition that may direct a preference of one target location to another.

**Fig 6.**
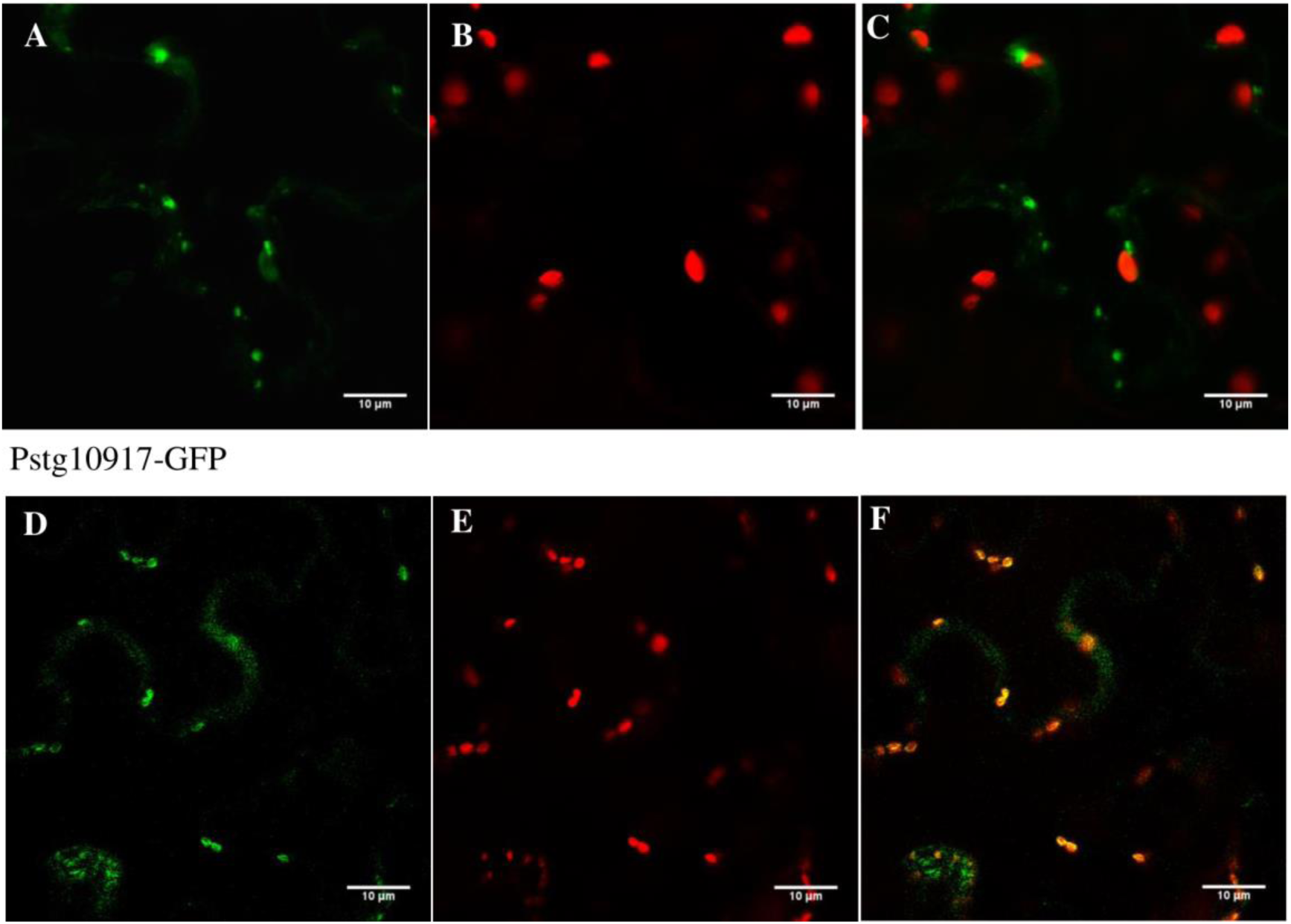
Subcellular localization of Unigene17495 (Pstg10917) tagged with GFP at C-terminus. (A) Expression of Pstg10917ΔSP-GFP (without N-terminus secretion signal). (B) and (E) Chloroplast autofluorescence. (C) and (F) Overlaid images. (D) Expression of Pstg10917 (with N-terminus secretion signal). The expression on *N. benthamiana* leaves was visualized under a confocal microscope at 2 dpi.

### Unigene17495 (Pstg10917) suppresses INF1-induced cell death

Unigene17495 (Pstg10917) is conserved among *Pst* relatives. However, none of the homologs have been previously studied for their function. Interestingly, an effector candidate of the soybean rust pathogen (*Phakopsora pachyrhizi*), PpEC82 [46], resembles Unigene17495 (Pstg_10917) with an e-value of e^−14^. PpEC82 was reported to localize to the cytosol and nucleus with strong aggregation [47] properties, unlike the chloroplast-targeting Unigene17495 (Pstg_10917) as we have demonstrated. Moreover, Qi *et al*. stated that PpEC82 was able to suppress BAX-induced cell death when expressed in yeast cells [47]. Correspondingly, we conducted cell death suppression assays to test whether Unigene17495 (Pstg_10917) has suppression abilities similar to its distant homolog, PpEC82. We used a cell death elicitor (INF1) as a cell death inducer and categorized the resulting suppression of cell death observations into four levels: strong suppression, moderate suppression, weak suppression, or no suppression at all. Across 15 replicates of Unigene17495-treated *N. benthamiana* leaves, we observed three cases of strong, three cases of moderate, and three cases of weak cell death suppression. The other six leaves presented no significant level of cell death suppression. In all 15 trials, leaves treated with the negative controls (GFP and SP-GFP) resulted in tissue collapse and cell death in the infiltrated area. Therefore, we concluded that Unigene17495 (Pstg_10917) without SP, was able to partially suppress programmed cell death triggered by the INF1 elicitor (Fig 7). We need to emphasize that not all attempts produced equal levels of cell death suppression. Although the expression of the effector was verified by microscope analysis at 2 dpi for all of the leaves, 6 out of the 15 leaves revealed no detectable suppression of INF1-induced cell death. Fisher’s exact test calculated a *P*-value of 0.0007 for the findings, and thus the results were significant (*P* < 0.05). Together, these data show that Pstg10917ΔSP-GFP (Unigene17495) is capable of significantly suppressing PCD induced by INF1.

**Fig 7.**
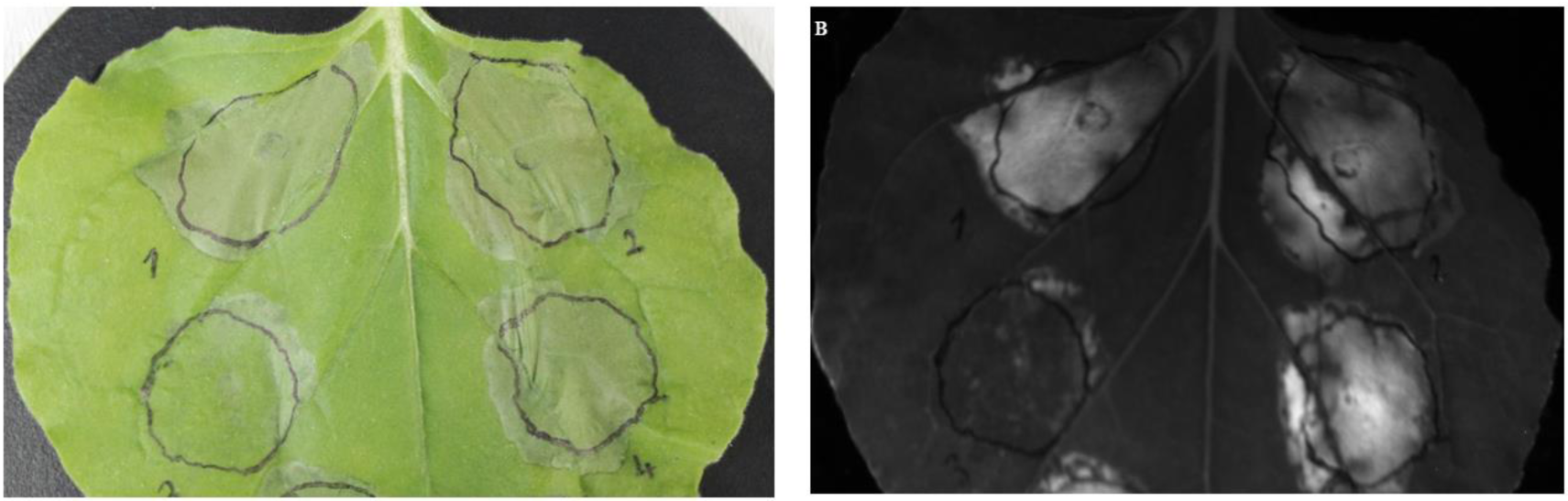
Suppression of cell death mediated by Inf1 expression. (A) Daylight and (B) UV light exposure. Constructs were expressed with *Agrobacterium tumefaciens*; (1) GFP, (2) SP-GFP, (3) Unigene17495 without SP (Pstg10917ΔSP-GFP) and (4) Unigene17495 (Pstg10917-GFP). Cell death elicitor (Inf1) was infiltrated into the same region after 24 hours. Photos were taken at 4 days after Inf1 challenge.

It should be noted that the complex nature of heterologous expression systems serves as a double-edged sword due to the oscillations in cell death response with *N. benthamiana*. However, the versatility of the system provides a fast and effective way to study candidate effectors. Other candidate effectors of *Pst* have been studied using heterologous expression systems. For example, the *Pst* effector PstHa5a23 was reported to suppress PCD triggered by BAX, INF1, and two mitogen-activated protein kinases (MKK1 and NPK1) involved in host cell resistance [48]. PstHa5a23 is a homolog to Pstg_00676 identified in our PstDESSPs list. In a recent publication, Pstg_14695 and Pstg_09266 were able to suppress PCD as part of effector-triggered immunity [49]. Furthermore, Petre *et al*. utilized mass spectroscopy analysis to decipher host cell factors that interact with GFP-tagged candidate effectors in co-immunoprecipitation samples [50]. Eleven of the twenty candidate effectors reported in that study are homologous to our PstDESSPs. These recent reports support the reliability of our PstDESSPs data as a trustworthy collection of candidate effectors that can be used for effector function studies in the future.

## Conclusion

The obligate biotrophic fungus *Pst* causes yellow rust disease by establishing an interface between the host and pathogen using small secreted proteins. Transcriptome sequencing has proven a useful strategy in elucidating the crosstalk of proteins between the host and pathogen, including the interaction between wheat and *Pst*. We compared compatible and incompatible interaction points in order to identify the differentially expressed genes (PstDEGs) at 10 dpi. *In silico* predictions highlighted the SSPs within the group of PstDEGs. The annotations, comparisons, and characterizations contributed to the identification of a list of the most promising subset of SSPs that function as phytopathogen effectors. The presented SSP data represent candidates for the functional analysis to better understand yellow rust disease in wheat. In particular, we were able to determine that Unigene17495 (Pstg10917) of our PstDESSPs list is localized to the chloroplasts of infected plant cells and is capable of halting programmed cell death.

## ACKNOWLEDGMENTS

This study was funded by COST-113Z350 (call FA1208), PI: MSA. BD (Dagvadorj), ACO, and AA-O were supported by TUBITAK-BIDEB: 2215 and 2211/C, respectively. The authors want to thank the Kamoun Lab, Sainsbury Lab, and Tolga Bozkurt of Imperial College London, Department of Life Sciences, London, SW7 2AZ, the UK for vectors used in the study.

## AUTHOR CONTRIBUTIONS

Ahmet Caglar Ozketen: Methodology, Investigation, Data Curation, Validation, Writing – original draft preparation, Writing – review & editing

Ayse Andac-Ozketen: Investigation, Validation

Bayantes Dagvadorj: Investigation

Burak Demiralay: Data Curation

Mahinur S. Akkaya: Methodology, Funding acquisition, Supervision, Writing – review & editing

## SUPPORTING INFORMATION

**Fig S1. Length distribution of contigs and unigenes after the assembly process.** Graphical representation of the length distribution of A) contigs of AvocetS_PST, B) unigenes of AvocetS_PST, C) contigs of AvocetYR10_PST and D) unigenes of AvocetS_PST.

**Fig S2. Function annotation of the unigene data.** The function of each of the unigenes was determined using the A) COG or B) GO databases.

**Table S1. Annotation of the assembled unigenes discovered in transcriptome sequencing.**

**Table S2. Annotation of the PstDEGs mapped to the *Pucciniales* local database.**

**Table S3. In-depth characterization of the differentially expressed small secretome profile of *Pst* (PstDESSPs).**

**Table S4. Primers used in this study for cloning.**

**Table S5. Matching PstDESSPs to the YR candidates identified in the study by Xia *et al*., 2017.**

