## Supplementary figures and images for "In-depth secretome analysis of *Puccinia striiformis* f. sp. *tritici* in infected wheat uncovers effector functions"

### Supplemental Figure 1

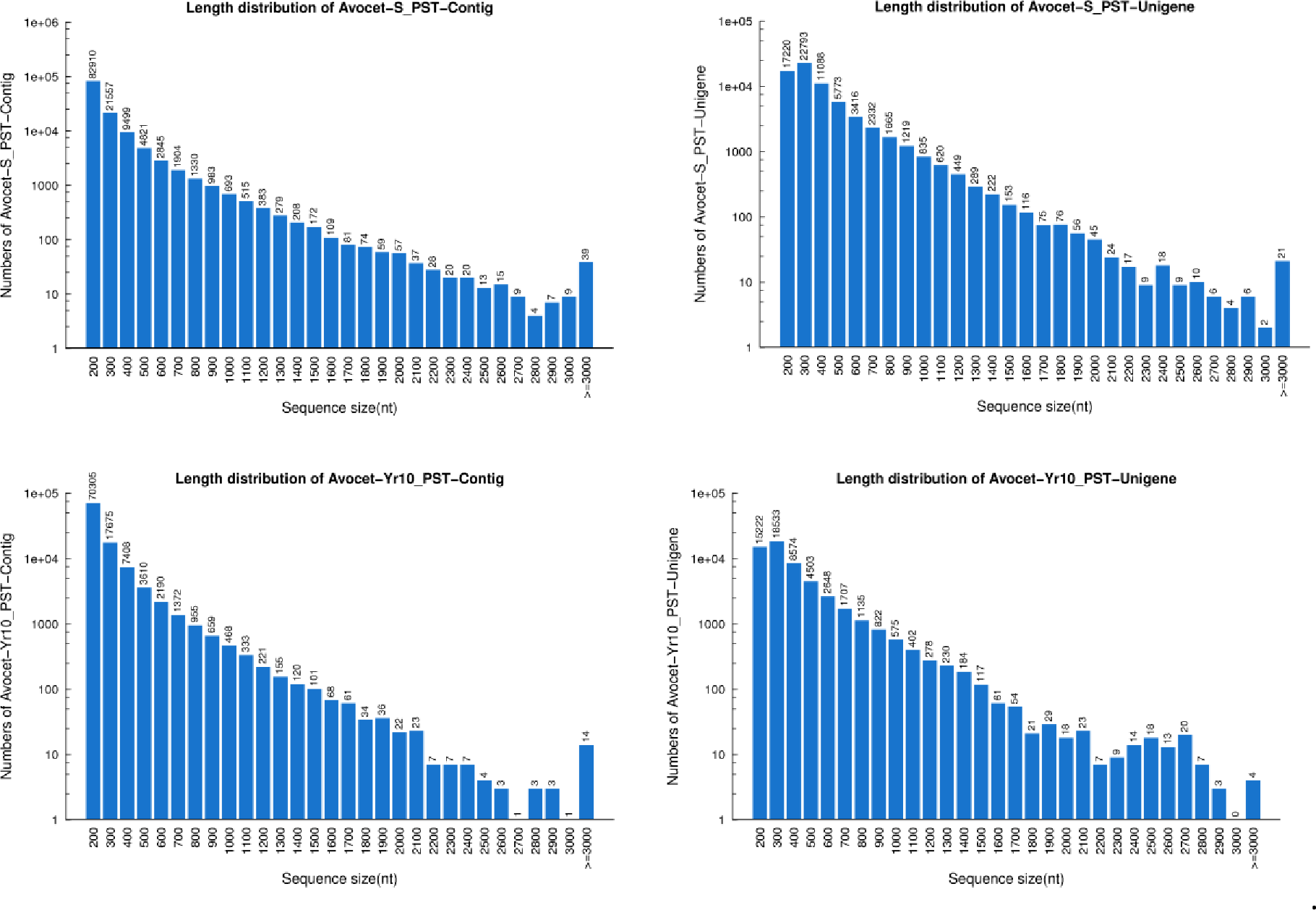

### Supplemental Figure 1

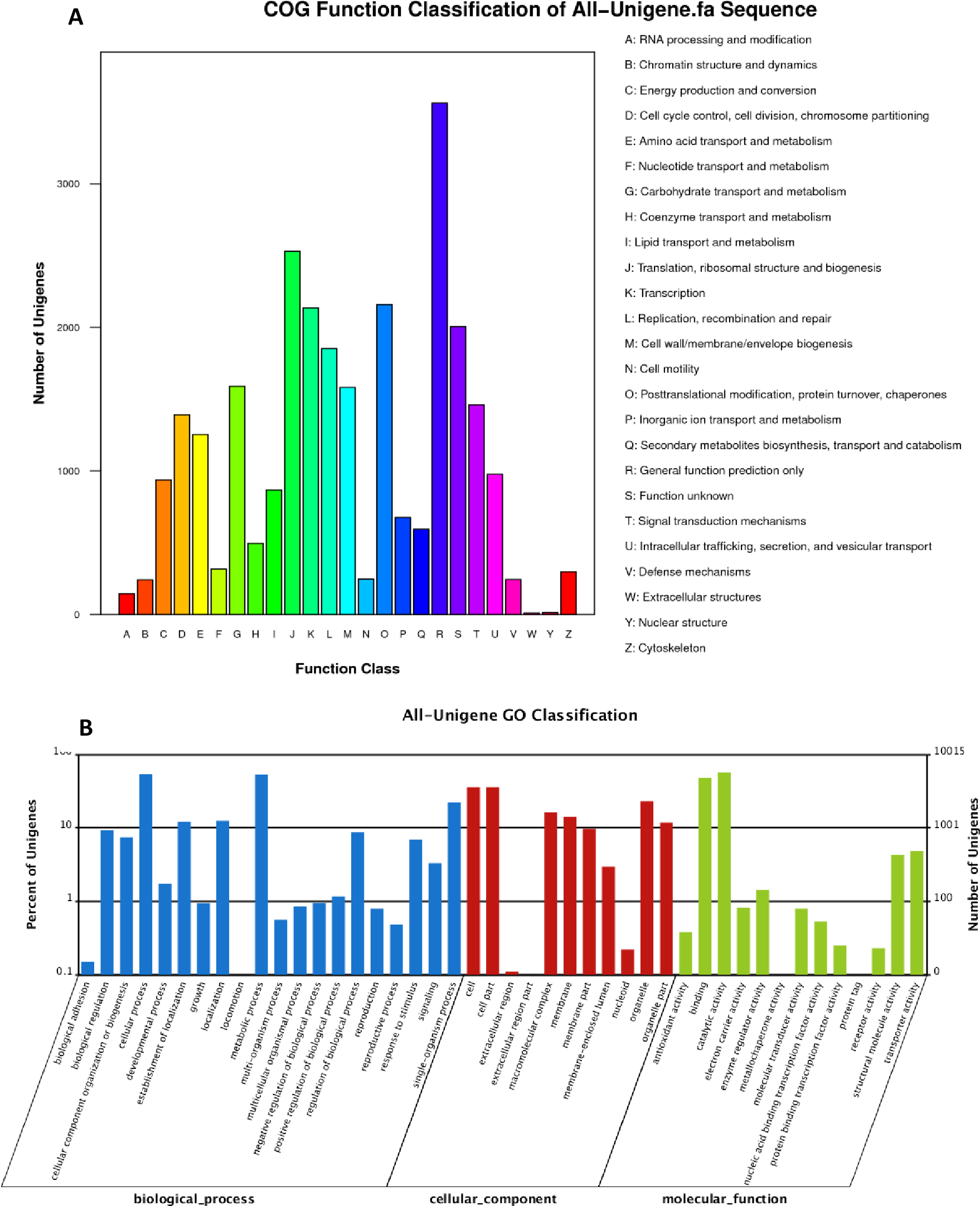
