## Supplemental Table 4 for "In-depth secretome analysis of *Puccinia striiformis* f. sp. *tritici* in infected wheat uncovers effector functions"

| Gene ID | Pucciniales homologs | Avr candidates |
| --- | --- | --- |
| CL3094.Contig2_All | PSTG_09464T0* | YR6 |
| CL6786.Contig1_All | PSTG_02003T0 | YR6 |
| Unigene30809_All | PSTG_03083T0 | YRTR1 |
| Unigene31162_All | PSTG_14206T0 | YR9 |
| Unigene31932_All | PSTG_08524T0 | YR9 |
| Unigene37241_All | PSTG_11923T0* | YR17 |
| Unigene37856_All | PSTG_16854T0* | YR9 |
| Unigene38264_All | PSTG_00708T0 | YR6 |
| Unigene7586_All | PSTG_14207T0 | YR9 |

**Table S4.** Matching PstDESSPs compared against YR candidates identified in Xia *et al*., 2017.

*Three of PstDESSPs overlapped with three previously neglected YR candidates in Xia et al., 2017.
