## Supplemental Table 5 for "In-depth secretome analysis of *Puccinia striiformis* f. sp. *tritici* in infected wheat uncovers effector functions"

| Primer Names | Sequences (5’-3’ direction) | Length (bp) |
| --- | --- | --- |
| CACC-SP-917F | CACCATGTTGTTCTACGTTTACCTCA | 26 |
| CACC-917F | CACCATGCAGACTTTACCTTCCG | 23 |
| 917Rev-STP | CTAGCATGTTTCCCAGCCTCC | 21 |
| 917Rev | GCATGTTTCCCAGCCTCCG | 19 |

**Table S5.** Primers used in this study for the cloning.
